# vGPCR-mediated ANO1 Modulation of Apoptosis Regulates Lytic KSHV Infection

**DOI:** 10.64898/2026.02.18.706543

**Authors:** Anisha Reddy Konakalla, Osvaldo K. Moreno, Savannah E. Price, Erica L. Sanchez

**Affiliations:** Department of Biological Sciences, The University of Texas at Dallas, Richardson, Texas, USA; Department of Molecular and Human Genetics, Baylor College of Medicine, Houston, TX, USA

## Abstract

Kaposi’s Sarcoma herpesvirus (KSHV) is an oncogenic, enveloped double-stranded DNA virus that causes Kaposi’s Sarcoma (KS), an endothelial-derived tumor that primarily affects immunocompromised individuals. KSHV persists in a biphasic lifecycle of latency and lytic reactivation, with the lytic phase driving viral dissemination and disease progression through expression of viral proteins that reprogram host survival pathways. One such protein is the viral G protein-coupled receptor (vGPCR), a constitutively active lytic gene product that promotes angiogenic and pro-survival signaling, although the host mediators of these effects remain incompletely defined. Here, we identify Anoctamin-1 (ANO1), a calcium-activated chloride channel with ant-apoptotic functions, as a downstream target of vGPCR in endothelial cells. RNA sequencing and RT-qPCR in vGPCR-expressing HMEC-1 cells revealed broad transcriptional reprogramming, including strong ANO1 induction. Functional studies showed that ANO1 depletion had little effect on viability in full-serum conditions but selectively sensitized vGPCR-expressing cells to apoptosis during serum starvation. This loss of viability was rescued by the pan-caspase inhibitor Z-VADFMK, indicating caspase-dependent apoptosis. To examine relevance during infection, we analyzed iSLK.BAC16 cells undergoing lytic reactivation. ANO1 was similarly upregulated during reactivation. Genetic knockdown or pharmacological inhibition of ANO1 with DES or Ani9 increased late-stage apoptosis while minimally affecting early apoptosis. Notably, ANO1 loss enhanced early and late lytic gene expression and increased production of infectious virions. These findings identify ANO1 as a vGPCR-induced host survival factor that suppresses apoptosis while modulating KSHV lytic replication, revealing a host ion-channel-dependent checkpoint linking cell survival to viral replication and KSHV pathogenesis.

**IMPORTANCE:** KSHV drives tumorigenesis by manipulating host cell survival pathways, yet how apoptosis is regulated during infection remains incompletely understood. We identify ANO1 as a host factor induced by the vGPCR that suppresses caspase-dependent apoptosis in endothelial cells. ANO1 is upregulated in both vGPCR-expressing endothelial cells and during KSHV lytic reactivation in iSLK.BAC16 cells. Disruption of ANO1 increases apoptotic cell death and is accompanied by elevated lytic gene expression and greater production of infectious virions, suggesting that apoptosis can coincide with enhanced lytic replication. These findings implicate host ion-channel signaling in the control of apoptosis during KSHV infection and indicate that the balance between cell survival and apoptosis influences viral gene expression and replication. Understanding how KSHV manipulates host survival pathways provides insight into mechanisms governing viral replication and may inform approaches to limit KSHV-associated disease.

## INTRODUCTION

Kaposi’s Sarcoma (KS), caused by Kaposi’s Sarcoma Herpesvirus (KSHV), is an endothelial cell-based tumor and the most common type of cancer in HIV-infected, untreated individuals [1,2]. KSHV is an oncogenic enveloped double-stranded DNA virus and belongs to the *Gammaherpesvirinae* subfamily. KSHV is also associated with lymphoproliferative disorders such as primary effusion lymphoma (PEL), Multicentric Castleman’s Disease (MCD), and KSHV inflammatory cytokine syndrome (KICS) [3]. KSHV infection of endothelial cells results in morphological, metabolic, lifespan, and gene expression changes to facilitate virion production.

KSHV pathogenesis exhibits two different modes of infection, latent infection and lytic reactivation. Most KS cells are in the latent phase by default, with only 1-5% lytically replicating [4,5]. The latent phase of KSHV is marked by limited viral gene expression, no virion production, and long-term host cell survival. However, during the lytic phase, all viral genes are expressed, and virion production occurs, which ultimately leads to host cell lysis [2]. Lytic viral proteins can drive oncogenesis, induce cellular proliferation, and modulate the host immune system [2]. The KSHV lytic cycle is initiated by expression of the immediate early gene and trans-regulator Replication and Transcriptional Activator (RTA, ORF50) [6].

Currently, the focus of KSHV research lies predominantly in the latent phase of infection, as latently infected cells comprise the majority of KS tumors and latency allows for long-term viral persistence and immune evasion [7]. However, the lytic replication stage remains underexplored, despite its critical role in viral dissemination, inflammation, and tumor progression. Many lytic genes are expressed during reactivation and act as potent modulators of host signaling pathways that promote various cellular processes of infected cells [8–10]. One such lytic gene is viral G Protein-Coupled Receptor (vGPCR), which is constitutively expressed during early reactivation and is known to be further activated by several cellular cytokines. vGPCR is a seven-transmembrane signaling protein, homologous to the cellular cytokine receptor CXCR2 [11–15]. Although extensive work has characterized oncogenic signaling pathways of vGPCR, far less is known about how vGPCR reshapes global host transcriptional programs in endothelial cells, particularly in the context of KSHV infection. Our approach to studying this gap involved examining endothelial cells overexpressing vGPCR to determine the downstream transcriptional regulation that is dependent on this viral transmembrane receptor. Understanding the mechanisms and impact of lytic gene expression can uncover new therapeutic vulnerabilities that are missed when focusing solely on latency.

Our RNA-sequencing data revealed several previously unknown putative vGPCR-targets that were upregulated, including Anoctamin-1 (ANO1), a transmembrane calcium chloride channel implicated in apoptosis inhibition, that also has a potential role in nucleotide regulation [16–18]. We investigated vGPCR-dependent upregulation of ANO1 to better understand the role this cellular protein plays in apoptosis. ANO1 has been shown to induce an anti-apoptotic effect via activation of the PI3K/Akt pathway in cancer models [18]. Interestingly, vGPCR has also been reported to induce an anti-apoptotic effect by activating the PI3K/Akt pathway when overexpressed [19]. However, a vGPCR-dependent regulation of ANO1 has not been reported. Our study addresses this gap by defining the relationship between vGPCR expression and ANO1 function and by evaluating how ANO1 contributes to apoptosis in our vGPCR overexpression system.

Building on these studies, we investigated the role of ANO1 in KSHV lytic replication and infectious virion production. We determined that ANO1, upregulated by the lytic viral gene vGPCR, regulates the production of infectious KSHV virions by modulating apoptosis. Using siRNA-mediated knockdown and pharmacological inhibition with DES and Ani9, we demonstrated that loss of ANO1 leads to increased apoptotic cell death. Interestingly, we also identified that inhibition of ANO1 significantly enhances the release of infectious virions from iSLK.BAC16 cells. These findings reveal that ANO1 functions as a key host factor limiting virion production, whereby its inhibition creates a cellular environment that facilitates more efficient KSHV replication and infectious virus release, highlighting a novel intersection between host ion channel activity and viral replication dynamics. Together, these findings reveal the importance of host ion channel regulation during KSHV infection, as understanding how ANO1 influences the balance between apoptosis and viral replication may reveal new therapeutic targets for controlling KSHV-associated pathogenesis.

## RESULTS

### vGPCR overexpression leads to global changes in the host cell transcriptome, including the upregulation of ANO1

Human microvascular endothelial cells (HMEC-1) were transfected with the vGPCR plasmid to assess the changes in host gene expression induced by vGPCR compared to the empty vector-transfected cells. A schematic representation of the experimental approach to determine gene expression changes in vGPCR-transfected endothelial cells is shown in **Figure 1A**. All samples were harvested 48 hours after transfection, followed by next-generation sequencing on mRNA libraries from empty vector (EV) and vGPCR samples in triplicate. Out of a total of 37,986 unique transcripts were successfully aligned, 49 annotated genes were differentially expressed (padj ≤ 0.05, fold change <-1 or ≥ 1) in empty control vs vGPCR-transfected cells as depicted in the volcano plot (**Figure 1B**). Our results show that 3 genes were downregulated, and 46 genes were upregulated, indicating that vGPCR expression and its downstream signaling induce a wide array of transcriptional changes in endothelial cells. Heatmap visualization of all the significantly differentially expressed genes across replicates further illustrated the consistent transcriptional reprogramming in vGPCR-overexpressing HMEC-1 cells **(Supplemental Figure 1A)**.

**Figure 1.**
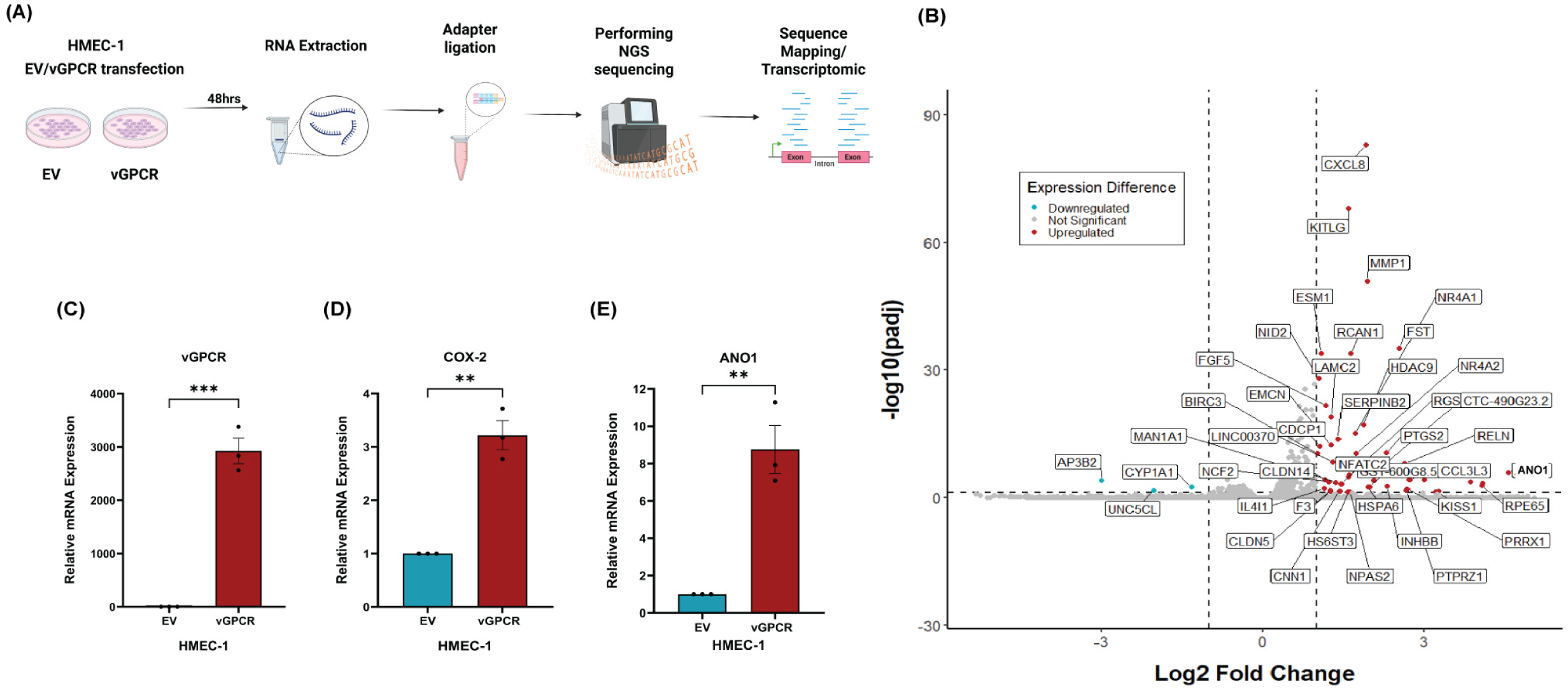
ANO1 is upregulated in vGPCR-expressing endothelial cells. **(A)** HMEC-1 Bulk RNA-sequencing workflow. HMEC-1 cells were either transfected with vGPCR or EV for 48 hours for bulk RNA-sequencing and RT-qPCR validation. **(B)** Volcano plot showing differentially expressed cellular transcripts in vGPCR-expressing cells compared to EV controls. **(C-E)** RT-qPCR gene expression validation in HMEC-1 cells expressing **(C)** the viral early lytic gene, vGPCR, and host genes **(D)** COX-2 and **(E)** ANO1 compared to EV controls. All expression data are relative to housekeeping gene expression (GAPDH). Generated in GraphPad. Data represent mean ± SEM from [n=3] independent experiments. Statistical significance was determined by t-test; ***, *P* ≤ 0.001; **, *P* ≤ 0.01. Genes with adjusted *p*-values 0.05, and Log2FoldChange -1, 1 were plotted. EV, Empty Vector.

Among the 46 upregulated genes, several were previously described to be increased in vGPCR-dependent host transcriptional alterations and genes related to the pro-inflammatory, pro-angiogenic pathways, viral lytic induction, and extracellular matrix remodeling. Cyclooxygenase 2 (COX2) [20,21], CXCL8 [14], KITLG [22], and RCAN1 [22] were all upregulated in our RNA-seq analysis **(Figure 1B**). Downregulated genes included CYP1A1, UNC5CL, and AP3B2. Gene ontology analysis of the top 10 significantly activated pathways revealed enrichment for pathways involved in cell movement and tissue remodeling, and the top 10 significantly suppressed pathways revealed downregulation of pathways associated with translational and ribosome biogenesis machinery, indicating suppression of global translational activity **(Supplemental Figure 1B**). The expression levels of some of the top upregulated genes in our HMEC-1 RNA-sequencing data, previously studied in vGPCR-dependent transcriptional alterations, including FST, CXCL8, HDAC9, and KITLG, were confirmed using RT-qPCR analysis **(Supplemental Figure 1C).**

To validate successful vGPCR transfection, we assessed mRNA expression of the canonical vGPCR target gene, COX-2 [20,21] in empty vector and vGPCR-transfected cells. RT-qPCR analysis confirmed that upon overexpression of vGPCR, COX-2 expression is significantly upregulated ∼2.5 fold compared to empty vector controls. (**Figure 1C and 1D)**. Furthermore, several previously unknown putative vGPCR-targets were identified to be upregulated, notably ANO1, a transmembrane calcium chloride channel implicated in apoptosis inhibition, cell proliferation, and tumor progression [16–18] **(Figure 1E)**. Studies show that ANO1 inhibits apoptosis and regulates the cell cycle [23]. We have thereby confirmed the RNA-sequencing data using RT-qPCR and observed that ANO1 was ∼8-fold upregulated in vGPCR-expressing endothelial cells. Taken together, these results provide strong evidence that the transcriptional changes in HMEC-1 cells are a direct consequence of vGPCR expression and its downstream signaling. Recent studies have shown that ANO1 can modulate host cell apoptosis through similar mechanisms as vGPCR [18,24]. Collectively, our data reveal that vGPCR drives broad transcriptional remodeling in endothelial cells and identifies ANO1 as a novel vGPCR- responsive gene that may help sustain the apoptosis-resistant phenotype in endothelial cells.

### Knockdown of ANO1 induces apoptotic cell death in HMEC-1 cells

Cell death can occur through several programmed or non-programmed mechanisms, including apoptosis (caspase-driven dismantling of cellular components), necroptosis (RIPK-mediated membrane rupture), autophagic cell death (excessive autophagosome formation), and pyroptosis (inflammasome-activated gasdermin pores) [18,24–27]. As previous studies have reported that ANO1 can modulate host cell apoptosis through mechanisms similar to those activated by vGPCR [28,29], we sought to test if vGPCR-mediated ANO1 upregulation modulates apoptotic cell death in endothelial cells. To investigate the functional role of ANO1 in vGPCR-mediated cell survival, we measured cell death percentages following ANO1 knockdown under both full serum and serum-starved conditions in HMEC-1 cells. Serum starvation is a well-established method to induce apoptotic cell death by depriving cells of external survival signals, including nutrients and growth factors. This approach is widely used to simulate stress conditions and to assess apoptosis sensitivity in endothelial cells, as previous studies have shown that serum deprivation reliably activates caspase-dependent apoptosis [30–33]. The knockdown efficiency of ANO1 in vGPCR-transfected HMEC-1 cells was observed to be ∼70% (**Supplementary Figure 3A**). Under full serum conditions, ANO1 knockdown did not significantly affect cell death in either empty vector or vGPCR-expressing cells, suggesting that ANO1 does not have a role in vGPCR-mediated survival under non-stressed conditions **(Figure 2A)**. In contrast, under serum-starved conditions, a significant difference was observed. vGPCR-expressing cells expressing ANO1 exhibited significantly lower cell death compared to both empty vector controls and vGPCR-expressing cells with ANO1 knockdown. Notably, cell death levels in vGPCR-expressing cells with ANO1 knockdown closely resembled those of serum-starved empty vector controls, indicating that the loss of ANO1 attenuates the protective effect conferred by vGPCR during nutrient deprivation. These results suggest that ANO1 is specifically required for vGPCR-mediated inhibition of apoptosis under serum-starved conditions, indicating ANO1 as a critical mediator of vGPCR-driven cell survival during stress.

**Figure 2.**
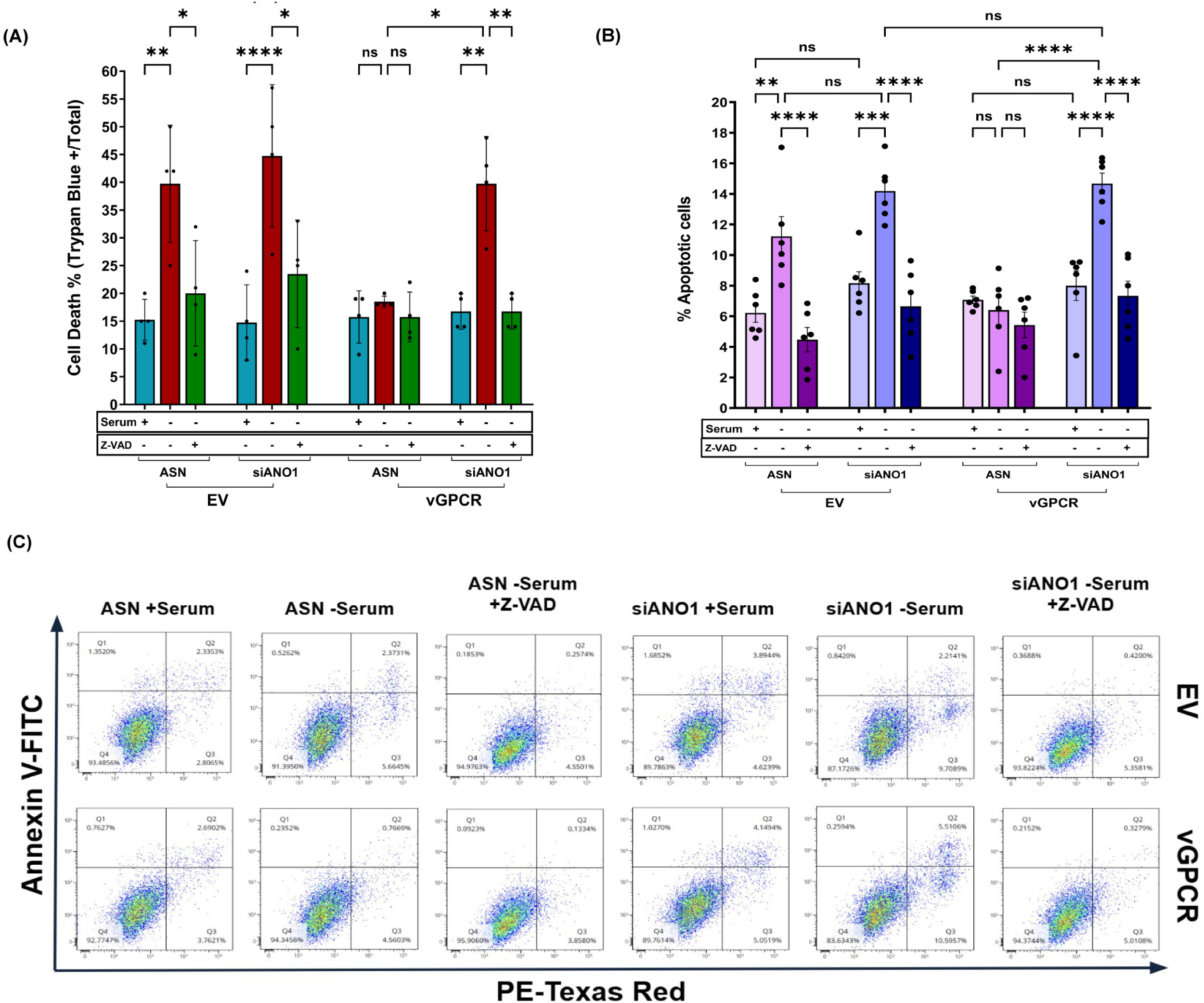
ANO1 knockdown induces caspase-dependent apoptotic cell death in vGPCR-expressing HMEC-1 cells under serum starvation. HMEC-1 cells transfected with vGPCR were treated with siANO1 and cultured under full serum or serum-starved conditions, with or without the pan-caspase inhibitor Z-VAD-FMK (25 µM). **(A)** Cell death was quantified by trypan blue exclusion assay in HMEC-1 cells transfected with EV or vGPCR and treated with control siRNA (ASN) or siANO1 under full serum (+) or serum-starved (–) conditions, in the presence or absence of the pan-caspase inhibitor Z-VAD-FMK. Serum starvation significantly increased cell death in EV cells and in vGPCR-expressing cells following ANO1 knockdown, which was partially rescued by Z-VAD-FMK treatment indicating that vGPCR-mediated ANO1 upregulation promotes cell death through an apoptotic mechanism. **(B)** Quantification of apoptotic cells by Annexin V-FITC/PI flow cytometry under the same conditions as in (A). Serum starvation increased the proportion of early-late and late apoptotic cells in EV controls, while vGPCR expression limited apoptosis. This protective effect was lost upon ANO1 knockdown and was significantly reversed by Z-VAD-FMK, indicating caspase-dependent apoptosis. **(C)** Representative Annexin V-FITC/PI flow cytometry plots showing viable (FITC⁻PI⁻), early apoptotic (FITC⁺PI⁻), and late apoptotic/dead (FITC⁺PI⁺ / FITC⁻PI⁺) populations in EV and vGPCR-expressing HMEC-1 cells under the indicated conditions. Generated in GraphPad. Data represent mean ± SEM from independent experiments (n ≥ 3). Statistical significance was determined by one-way ANOVA; ****, *P* ≤ 0.0001; ***, *P* ≤ 0.001; **, *P* ≤ 0.01; *, *P* ≤ 0.05; ns, non-significant; EV, Empty Vector; ASN=All-Star Negative.

To determine whether the observed cell death upon ANO1 knockdown in vGPCR-expressing cells is due to apoptosis, we performed cell viability assays in the presence or absence of the pan-caspase inhibitor Z-VAD-FMK. vGPCR-transfected HMEC-1 cells treated with siANO1 showed a significant increase in cell viability when treated with Z-VAD-FMK compared to DMSO-treated controls **(Figure 2A)**, indicating that the loss of viability was caspase-dependent. This apoptosis rescue effect confirms that ANO1 knockdown induces apoptosis of vGPCR-expressing cells and further supports the role of ANO1 in mediating vGPCR-driven inhibition of apoptosis in endothelial cells. These results demonstrate that ANO1 is a critical effector of vGPCR-mediated cell survival under stress conditions. Although ANO1 is not required for vGPCR-mediated cell-survival under nutrient-rich conditions, it becomes essential for vGPCR to inhibit apoptotic cell death during serum starvation. The caspase-dependent nature of cell death following ANO1 knockdown further confirms its role in suppressing apoptosis. These results support a model in which vGPCR upregulates ANO1 to promote endothelial cell survival during lytic KSHV infection, highlighting ANO1 as a key mediator of vGPCR-driven anti-apoptotic signaling.

To more precisely quantify apoptosis, we repeated the serum starvation experiment by flow cytometry using Annexin V-FITC and PI staining. Consistent with our viability assays, ANO1 knockdown had no effect under full serum conditions, with both EV and vGPCR-expressing cells showing high viability, with ∼90% cells remaining viable (FITC^-^ PI^-^, Q4) regardless of ANO1 knockdown while early-late + late apoptotic cells (FITC^+^PI^+^ + FITC^-^PI^+^) remained low (<10%), confirming that ANO1 loss does not impact survival under nutrient-rich conditions **(Figure 2B, 2C)**. However, serum starvation revealed a clear requirement for ANO1 in vGPCR-mediated survival. Serum starvation substantially increased early-late and late apoptosis in EV cells, with the proportion of FITC^+^PI^+^ + FITC^-^PI^+^ cells rising from 6-8% in full serum to ∼12-15% in serum-starved cells. vGPCR expression partially protected against this increase, maintaining most cells in Q4 (>90%) and limiting apoptosis to <10%. While vGPCR expression reduced early-late and late apoptotic cells (FITC^+^PI^+^ + FITC^-^ PI^+^) compared to EV controls, this protective effect was lost when ANO1 was silenced, resulting in a marked increase in this apoptotic cell population, from <10% in full serum to ∼15% in serum-starved cells. Notably, apoptosis levels in vGPCR + siANO1 cells closely resembled those of serum-starved EV control. Similar to the trypan blue viability assay, treatment with the pan-caspase inhibitor Z-VAD-FMK significantly reduced the proportion of early-late and late apoptotic cells (FITC^+^PI^+^ + FITC^-^ PI^+^) in serum-starved vGPCR-expressing cells following ANO1 knockdown, with a corresponding increase in viable cells (FITC^-^PI^-^), indicating caspase-dependent apoptosis. These data confirm that ANO1 is specifically required for vGPCR-dependent inhibition of apoptosis during nutrient stress, consistent with its proposed role as a key mediator of vGPCR-driven anti-apoptotic signaling.

### Global transcriptomic changes in iSLK.BAC16 cells during KSHV lytic reactivation

To investigate the involvement of host factors in KSHV viral gene expression, we utilized iSLK.BAC16 cells, a well-established cell line that stably carries the BAC16 KSHV genome and expresses the viral lytic switch protein RTA (Replication and Transcriptional Activator) under the control of a doxycycline-inducible promoter, and encodes a constitutively expressed GFP reporter [34]. To assess host gene expression during lytic KSHV infection, we measured the expression of ANO1 in iSLK.BAC16 cells with and without the induction of lytic reactivation by RT-qPCR analysis. A schematic representation of the experimental approach to determine gene expression changes in reactivated iSLK.BAC16 compared to the non-reactivated iSLK.BAC16 is shown in **Figure 3A**. iSLK.BAC16 cells were reactivated using doxycycline (Dox) and sodium butyrate (NaB) for 48 hours, and RNA was extracted for next-generation sequencing (RNA-sequencing). Gene expression profiles were compared between uninduced (latent) and reactivated (lytic) conditions to identify differentially expressed genes. Out of 47,403 unique transcripts aligned, 3,269 genes were found to be significantly differentially expressed (padj < 0.005 and log2FoldChange -2, 2). Among these, 3,056 genes were upregulated, and 213 were downregulated. Volcano plots visualizing differential gene expression reveal a clear shift toward upregulation of host transcripts during lytic reactivation, suggesting a substantial remodeling of the host transcriptome in response to KSHV lytic gene expression **(Figure 3B)**. Importantly, several of the genes found to be upregulated in vGPCR overexpressing HMEC-1 cells were also observed to be elevated during KSHV reactivation in iSLK.BAC16 cells. These include HSPA6, PTGS2, RGS7, NCF2, NPAS2, INHBB, HDAC9, PTPRZ1, NR4A1, NID2, IL4I1, ANO1 and HS6ST3. These genes are involved in pathways involving angiogenesis, epithelial cell adhesion, IL18 signaling, and cancer. This convergence reinforces the relevance of the HMEC-1 vGPCR model for studying host responses to KSHV signaling.

**Figure 3.**
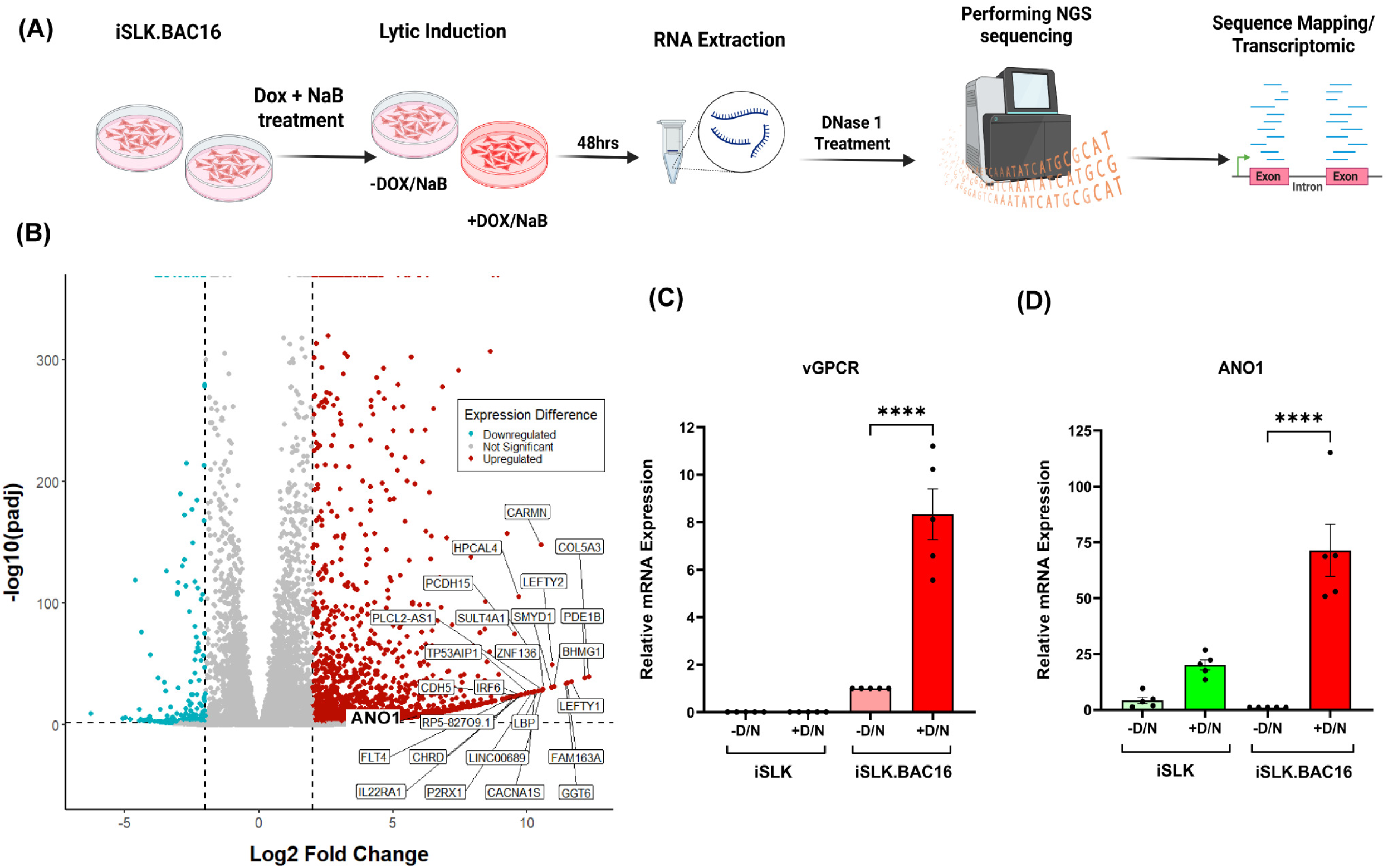
ANO1 is upregulated in iSLK.BAC16 cells during lytic reactivation. **(A)** Bulk RNA-sequencing experimental setup. **(B)** Volcano plot showing differentially expressed cellular transcripts in iSLK.BAC16 cells in the presence or absence of Dox and NaB. Only annotated genes are shown in the volcano plot. Relative mRNA expression of **(C)** viral gene vGPCR and **(D)** host gene ANO1 in lytic iSLK.BAC16 cells compared to the latent controls. Expression levels were normalized to GAPDH and compared between reactivated (48 h D/N treatment) and non-reactivated controls. Generated in GraphPad. Data represent mean ± SEM from [n=5] independent experiments. Statistical significance was determined by one-way ANOVA; ****, *P* ≤ 0.0001. D/N=Doxycycline/Sodium Butyrate. Genes with adjusted p values 0.05, and Log2FoldChange -2, 2.

As a confirmation that cellular mRNAs were differentially expressed during latency and following lytic reactivation, we performed RT-qPCR analysis using iSLK.BAC16 cells and compared them to a well-established, latently infected, RTA-inducible SLK cell line (iSLK) [35]. We observed that the expression levels of vGPCR were significantly upregulated in iSLK.BAC16 cells following lytic induction with ∼7-fold increase compared to non-reactivated iSLK.BAC16 cells and iSLKs **(Figure 3C).** Similarly, expression levels of the host gene ANO1 was also significantly upregulated with ∼70-fold increase, compared to non-reactivated iSLK.BAC16 cells. Notably, Dox and NaB alone upregulated the expression level of ANO1 by ∼20-fold **(Figure 3D).** The upregulation of ANO1 in reactivated iSLK.BAC16 cells is similar to our observations in HMEC-1 cells. ANO1 expression was among the most significantly upregulated genes in the lytic iSLK.BAC16 transcriptome (Log2FoldChange 5.984), suggesting that KSHV may exploit ANO1-mediated pathways during reactivation to support host cell survival or modulate ion homeostasis.

In addition, FABP4 and FABP7, members of the fatty acid-binding protein family involved in lipid metabolism and inflammation, were also upregulated, suggesting a potential link between lipid metabolic reprogramming and viral reactivation. Metabolism-associated pathways were prominently enriched among the upregulated genes, including fatty acid biosynthesis, glycolysis/gluconeogenesis, and oxidative phosphorylation [36,37]. This reprogramming is consistent with the metabolic demands of viral replication and may reflect a host attempt to control cellular homeostasis under stress. Several genes previously reported to be modulated during KSHV lytic replication were also differentially expressed in our dataset, including PTGS2 [38–40], RUNX3 [41], ISG15 [42], BATF3, IRF4, IRF6 [43], TLR5 [44], CXCR4 [45], SOCS3 [46,47], JAK2 [2,48], STAT5A [49]. These genes are known to participate in inflammatory signaling, oxidative stress response, and cell survival pathways.

Heatmap visualization of the top 50 upregulated genes across replicates further illustrated the consistent transcriptional reprogramming in reactivated iSLK.BAC16 cells **(Supplementary Figure 2A).** ANO1 was ranked 411th most upregulated gene among the 3,056 total genes upregulated in our differential expression analysis (out of only the significantly upregulated genes; ones with non-significant p-values were not included in the ranking). Gene ontology analysis of the top 10 most significantly suppressed pathways and the top 10 most significantly activated pathways is shown in **Supplementary Figure 2B.** Differentially expressed genes in latent versus lytic iSLK.BAC16 cells were significantly enriched in GO terms associated with muscle system processes, extracellular matrix components, calcium signaling, and immune regulation, suggesting a coordinated response involving cell morphogenesis, matrix remodeling, and apoptosis-related immune activation. Taken together, our transcriptomic analysis of iSLK.BAC16 cells following KSHV reactivation reveal a robust upregulation of host genes that mirror vGPCR-driven transcriptional programs, including inflammatory cytokines [14,50,51], pro-survival factors [21,52,53], and metabolic regulators [54,55]. These results provide further evidence that vGPCR contributes to shaping the host response during lytic replication and that targets such as ANO1 may serve as novel mediators of KSHV pathogenesis.

### Knockdown and pharmacological inhibition of ANO1 induce apoptosis during KSHV infection

Because our earlier findings demonstrated that ANO1 promotes survival of vGPCR-expressing endothelial cells under stress, we next asked whether ANO1 regulates apoptosis during KSHV infection. ANO1 has also been implicated in regulating cell survival pathways in cancer and viral infection contexts [18,24]. To test whether ANO1 influences apoptotic responses during KSHV reactivation, we performed siRNA-mediated knockdown of ANO1 in iSLK.BAC16 cells, followed by lytic induction with Dox and NaB. The knockdown efficiency of ANO1 in BAC16 cells was observed to be ∼75% **(Supplementary Figure 3B).**

We measured apoptosis by flow cytometry using Annexin V-APC and PI staining, allowing discrimination of early and late apoptotic populations. In cells transfected with siANO1, at 48 hours post-reactivation, no significant difference was observed in early apoptotic populations compared to ASN control cells under either latent or lytic conditions **(Figure 4A)**. However, late apoptotic cells increased ∼1.5 fold upon ANO1 knockdown following lytic induction compared to the ASN control, indicating that ANO1 primarily functions during lytic replication to limit cell death. Notably, Dox alone led to a ∼1-fold increase in late apoptotic cells following lytic induction compared to latent controls.

**Figure 4.**
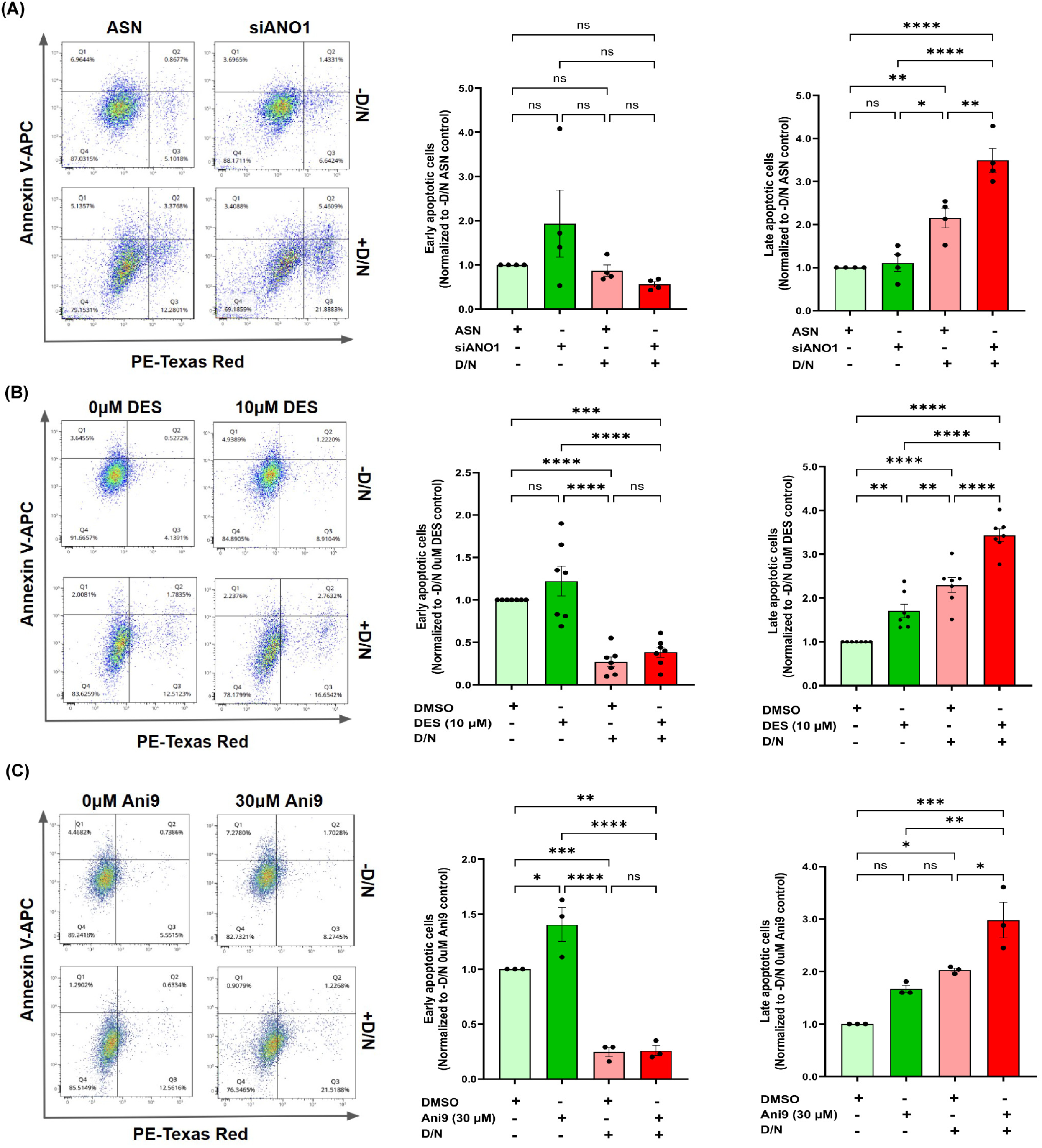
siRNA knockdown or pharmacological inhibition of ANO1 increases late apoptosis during KSHV lytic reactivation. iSLK.BAC16 cells were transfected with siANO1 or treated with the ANO1 inhibitors diethylstilbestrol (DES, 10 μM) or Ani9 (30 μM), followed by lytic induction with doxycycline and sodium butyrate (D/N) for 48 hours. Apoptosis was measured by flow cytometry using Annexin V-APC and Propidium Iodide (PI) staining to distinguish early and late apoptotic populations. A significant increase in late apoptotic cells in reactivated lytic samples but not in latent cells were measured upon **(A)** siRNA-mediated knockdown of ANO1 **(B)** DES treatment and **(C)** Ani9 treatment. , While a significant increase in late apoptotic cells was measured, early apoptotic cells remained unchanged in all conditions. Bars represent mean ± SEM from [n ≥ 3] independent experiments; individual biological replicates are shown as dots. Statistical comparisons were performed using one-way ANOVA with multiple comparison correction (****, *P* ≤ 0.0001; ***, *P* ≤ 0.001; **, *P* ≤ 0.01; *p ≤ 0.05; ns, not significant). D/N=Doxycycline/Sodium Butyrate; ASN=All-Star Negative.

Pharmacological inhibition of ANO1 using selective inhibitors DES (diethylstilbestrol) or Ani9 has been reported in connection with apoptosis regulation [56,57]. DES is a selective ANO1 inhibitor that induces apoptosis in various cancer cells by increasing caspase-3 activity and PARP-1 cleavage, with effective concentrations generally around 10 μM, showing anticancer effects without hepatotoxicity at this dose. DES has been previously shown to reduce both ANO1 channel activity and cell viability, reduce p-ERK1/2 and p-EGFR levels, and induce apoptosis by increasing caspase-3 activity and PARP-1 cleavage; however, it does not affect ATP-induced increase in intracellular Ca2^+^ levels [56]. Ani9 is a potent and highly selective small-molecule ANO1 inhibitor with an IC50 below 3 μM, previously used at concentrations ranging from 10 to 30 μM to robustly block ANO1 channel activity and induce apoptosis in prostate cancer cells, without affecting related chloride channels or intracellular calcium signaling [56–60]. These inhibitors have been consistently shown to increase apoptosis in ANO1-expressing cells, supporting their authenticity and functional relevance in studying the role of ANO1 in cell survival and death. DES and Ani9 treatment aligns with previous research demonstrating efficacy in inducing apoptosis through ANO1 inhibition, which complements the genetic knockdown approach by confirming the role of ANO1 in modulating apoptotic pathways.

iSLK.BAC16 cells were treated with the selective ANO1 inhibitors DES (10 μM) and Ani9 (30 μM) in the presence or absence of Dox and NaB induction of reactivation. Both drugs significantly increased apoptosis compared to vehicle controls, with reactivated cells showing a higher apoptotic rate than untreated or latent cells. Upon DES treatment, a significant increase in late apoptotic cells was observed under both latent (∼0.5-fold increase) and lytic (∼1.5-fold increase) conditions; however, the effect was markedly more pronounced during lytic reactivation **(Figure 4B)**. This suggests that ANO1 inhibition by DES compromises cell survival even in the absence of reactivation stimuli, but the apoptotic response is further amplified once lytic gene expression is induced. Similarly, Ani9 treatment also led to elevated late apoptotic populations upon reactivation (∼1-fold increase) while the early apoptotic cells remained unchanged during lytic reactivation, confirming that disruption of ANO1 function enhances apoptotic cell death during KSHV reactivation **(Figure 4C)**. This suggests that ANO1 inhibition induces a significant increase in late apoptosis of cells undergoing lytic reactivation. Our flow cytometry analysis also revealed that Dox and NaB treatment alone modestly increased the proportion of late apoptotic cells during the knockdown or pharmacological inhibition of ANO1, consistent with apoptosis associated with viral reactivation.

### ANO1 inhibition promotes early and late KSHV lytic gene expression

To determine whether ANO1 regulates the production of infectious KSHV virions, we knocked down ANO1 expression in iSLK.BAC16 cells using siRNA, or pharmacological inhibition via treatment with Ani9 or DES. RNA was collected after 48 hours of reactivation in the presence or absence of ANO1 inhibition, and RT-qPCR was performed to determine the gene expression levels of KSHV lytic genes. Upon reactivation, early lytic genes vGPCR and ORF59 relative mRNA expression was significantly increased by ∼0.5-fold and ∼1.5-fold, respectively, after the knockdown of ANO1 compared to the ASN control **(Figure 5A).** Similarly, the late lytic genes K8.1 and ORF26 were significantly upregulated by ∼2.5 fold and ∼1.5 fold, respectively, in the siANO1 knocked-down cells compared to the siASN control, indicating enhanced lytic reactivation.

**Figure 5.**
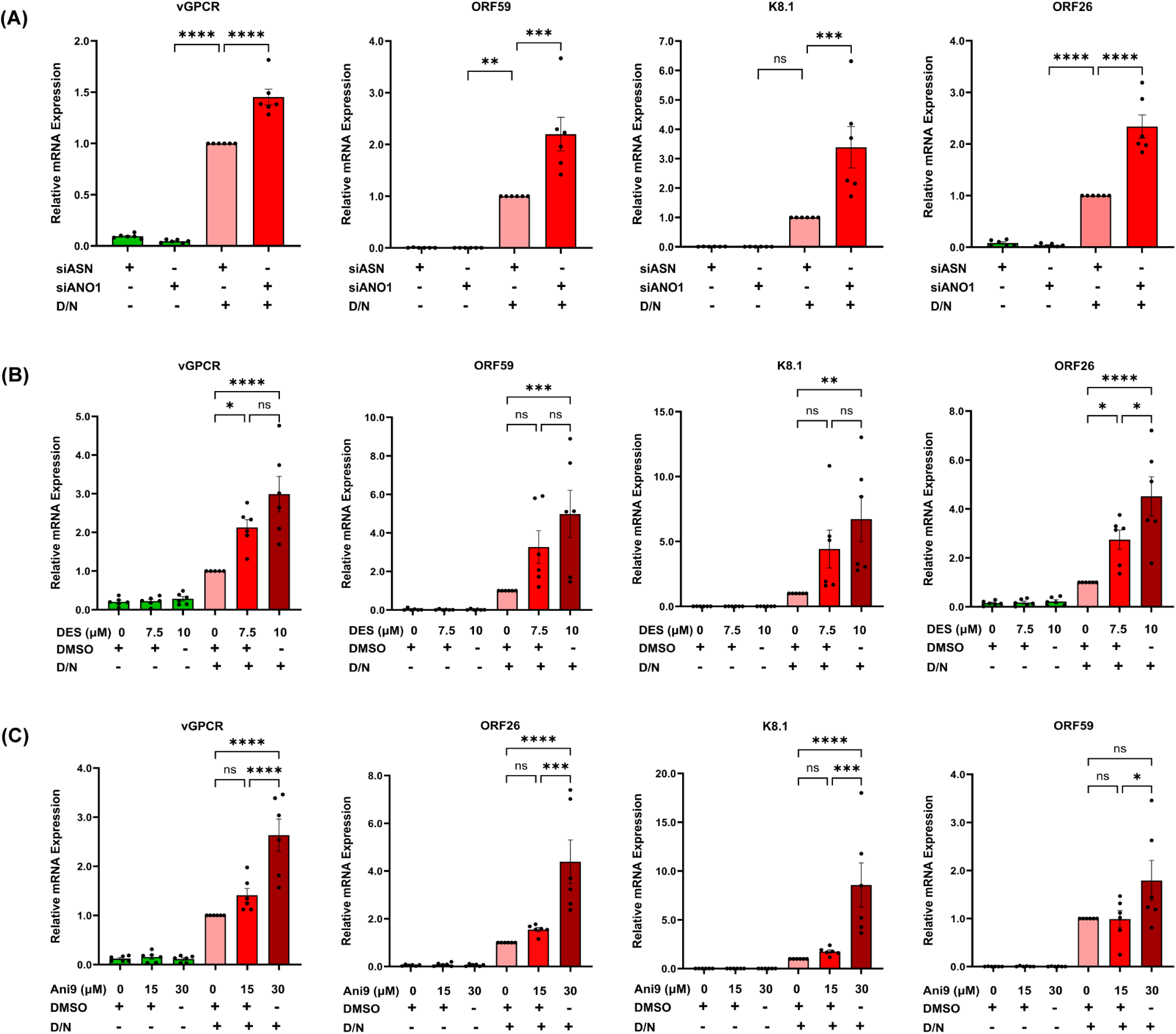
siRNA knockdown or pharmacological inhibition of ANO1 leads to increased viral lytic gene expression. iSLK.BAC16 cells were transfected with siANO1 or treated with Ano1 inhibitors diethylstilbestrol (DES, 10 μM) or Ani9 (30 μM), followed by lytic induction with doxycycline and sodium butyrate (D/N) for 48 hours. Relative mRNA expression of early lytic genes, i. vGPCR ii. ORF59 and late lytic genes, iii. K8.1 iv. ORF26 was measured by RT- qPCR. Viral gene expression was significantly upregulated upon **(A)** ANO1 knockdown, **(B)** Ani9 treatment, and **(C)** DES treatment, lytic reactivated iSLK.BAC16 cells. All expression data are relative to the expression of the housekeeping gene Tubulin. Generated in GraphPad. Data represent mean ± SEM from [n=6] independent experiments. Statistical significance was determined by one-way ANOVA; ****, *P* ≤ 0.0001; ***, *P* ≤ 0.001; **, *P* ≤ 0.01; *p ≤ 0.05; ns, non-significant. D/N=Doxycycline/Sodium Butyrate; ASN=All-Star Negative.

Treatment with Ani9 and DES resulted in a dose-dependent increase in both early and late lytic gene expression. Treatment with lower concentrations of DES (7.5 μM) resulted in non-significant increase in lytic gene expression, notably, higher doses (10 μM DES and 30 μM Ani9) resulted in significant increase in both early lytic, vGPCR (∼2 fold increase) and ORF59 (∼4 fold increase), and late lytic, K8.1 (∼6.5 fold increase) and ORF26 (∼4 fold increase) gene expression compared to the no drug control **(Figure 5B)**. Similarly, treatment with lower concentrations of Ani9 (15 μM) resulted in non-significant increase in lytic gene expression, but, higher doses (30 μM Ani9) resulted in significant increase in both early lytic, vGPCR (∼1.5 fold increase) and ORF59 (∼0.5 fold increase), and late lytic, K8.1 (∼7 fold increase) and ORF26 (∼3.5 fold increase) gene expression compared to the no drug control **(Figure 5C)**. These results suggest a dose-dependent effect of ANO1 inhibition on KSHV reactivation. Strikingly, all three ANO1-targeting approaches, siANO1, DES, and Ani9, produced the same outcome: a marked increase in the early and late lytic gene expression. Together, these findings indicate that ANO1 acts as a modulator of lytic gene expression, and its inhibition potentially promotes viral reactivation.

### ANO1 inhibition enhances the production of infectious KSHV particles

To assess whether the host gene, ANO1, influences the release of infectious KSHV particles, we measured viral titers following ANO1 knockdown and pharmacological inhibition during lytic reactivation. We performed siRNA-mediated knockdown/inhibition of ANO1 in iSLK.BAC16 cells followed by lytic reactivation. We collected the cell-free KSHV-containing supernatants from siANO1 and drug-treated samples and titered them on iSLK cells. We determined the infection levels in iSLK cells by quantification of GFP+ cells via fluorescence microscopy. Our results show a significant increase in the production of infectious virions, as determined by the elevated percentage of infection-after knockdown of ANO1, with ∼300% increase **(Figure 6A)** in infection compared to the siASN; ∼500% increase after inhibition with DES **(Figure 6B)**, and ∼800% increase after inhibition with Ani9 **(Figure 6C)** compared to no drug control. Notably, DES and Ani9 treatments did not result in an increase in viral titers at lower concentrations (7.5 μM DES and 15 μM Ani9), but significantly increased the infectious particles with higher concentrations of the drugs (10 μM DES and 30 μM Ani9).

**Figure 6.**
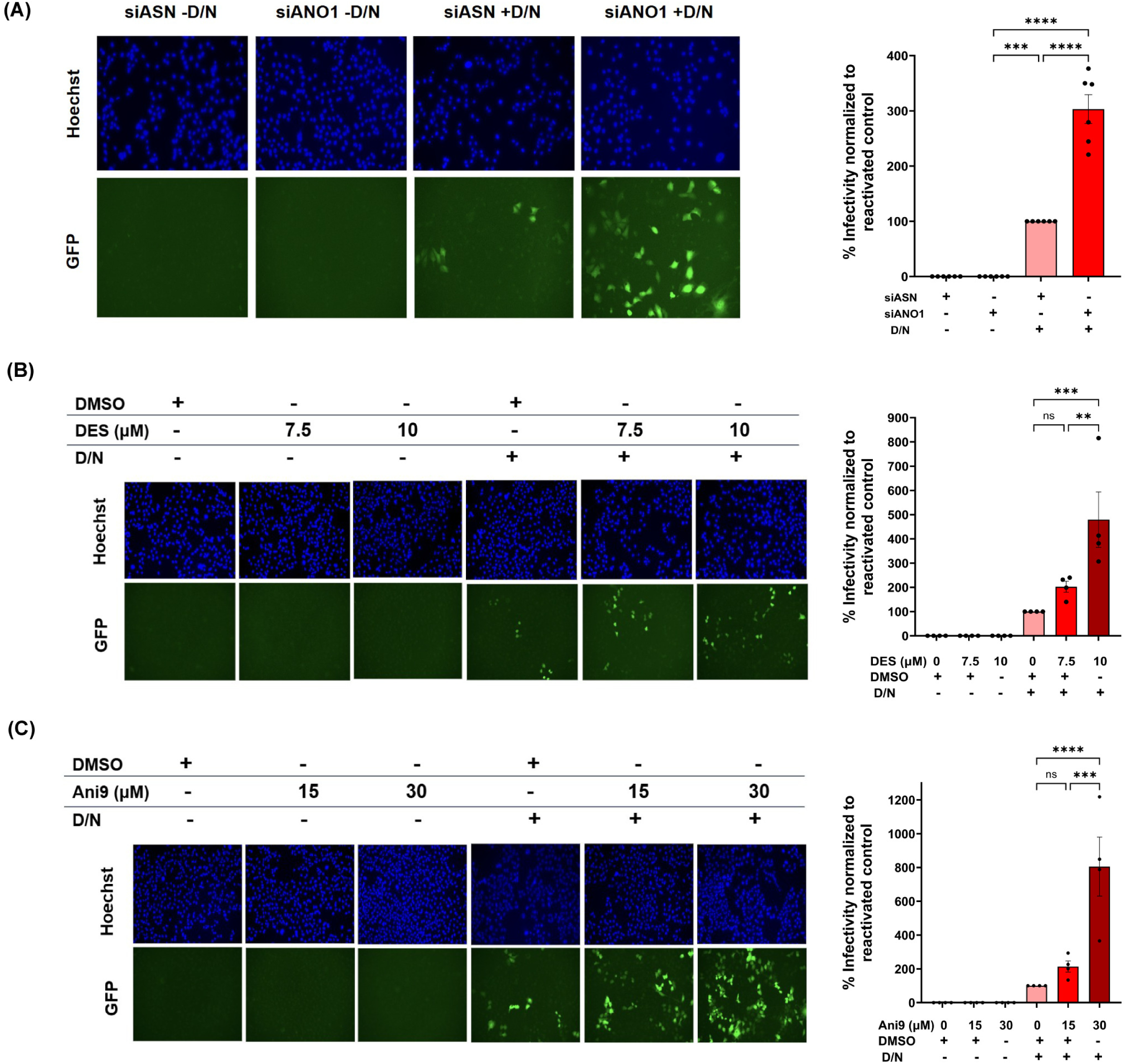
siRNA knockdown or pharmacological inhibition of ANO1 leads to increased extracellular viral titers. Cell-free supernatants from iSLK.BAC16 cells after knockdown or inhibition of ANO1 were collected after reactivation, and viral titers were determined. GFP+ cells correspond to KSHV-infected cells. Titers were assessed at 48 hpi. on a ZOE fluorescent microscope at 10X. **(A)** Lytic reactivated, siRNA-mediated knockdown of ANO1 significantly increased the production of infectious extracellular KSHV particles when titered on iSLK cells. Quantification of GFP+ cells observed in (A). **(B & C)** Pharmacological inhibition of ANO1 using Ani9 **(B)** and DES **(C)** resulted in a dose-dependent increase in extracellular viral titers, with higher concentrations promoting a significant increase in viral release. Imaged using ZOE, ImageJ was used to quantify the relative % of infectivity to reactivated control (siASN +D/N or DMSO +D/N). Generated in GraphPad. Data represent mean ± SEM from [n ≥ 3] independent experiments. Statistical significance was determined by one-way ANOVA; ****, *P* ≤ 0.0001; ***, *P* ≤ 0.001; **, *P* ≤ 0.01; ns, non-significant. D/N=Doxycycline/Sodium Butyrate; ASN=All-Star Negative.

**Figure 7.**
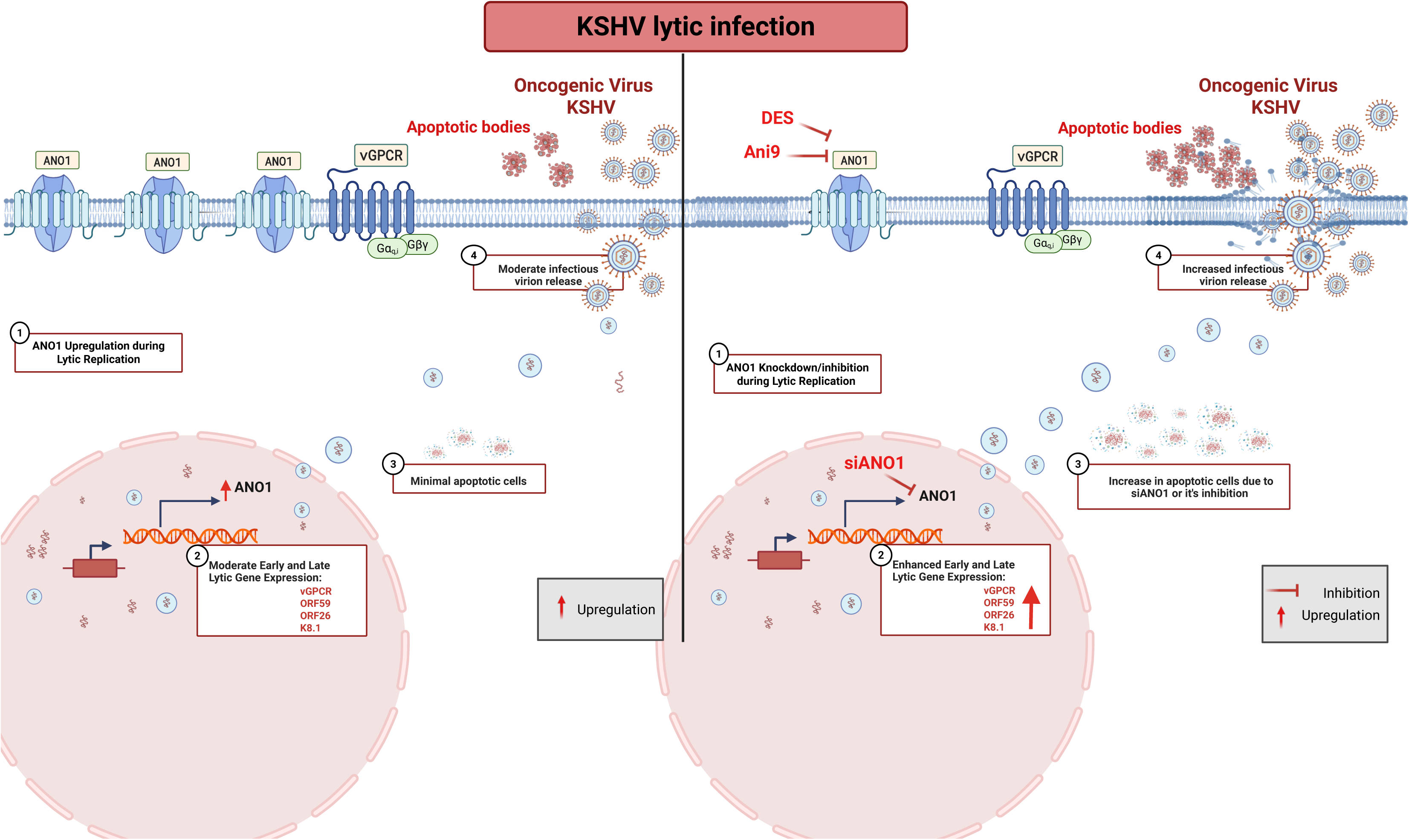
ANO1 modulates apoptosis and virion release during KSHV lytic infection. During KSHV infection, expression of the vGPCR is induced in the lytic phase, leading to transcriptional upregulation of the host calcium-activated chloride channel ANO1. Elevated ANO1 expression helps maintain cell survival by suppressing apoptosis, thereby supporting sustained viral replication within infected cells. In contrast, inhibition or depletion of ANO1 either by siRNA mediated knockdown or pharmacological inhibitors DES and Ani9 disrupts this protective effect, resulting in enhanced apoptotic cell death. Interestingly, this increased apoptosis coincides with a significant rise in the release of infectious KSHV virions, suggesting that ANO1 acts as a host restriction factor that limits virion egress during lytic replication. Collectively, these findings reveal that ANO1 functions as a key regulator of the cellular environment during KSHV lytic infection, balancing apoptosis and viral release to influence the efficiency of KSHV replication.

As convergent inhibition of ANO1 by three independent methods consistently enhanced early and late gene expression **(Figure 5)** and increased infectious virion release **(Figure 6)**, these findings suggest that ANO1 promotes infectious virion production, enhancing KSHV lytic replication and the release of infectious virus. Given our earlier observations that ANO1 loss promotes late apoptosis during KSHV infection **(Figure 4)**, the increase in extracellular virus titers upon ANO1 inhibition may be mechanistically linked to apoptotic membrane remodeling or vesicle release pathways that facilitate viral release, suggesting that cellular stress and ANO1 dysfunction create a permissive environment for productive viral release.

## DISCUSSION

Here, we demonstrate that vGPCR expression in endothelial cells induces transcriptional reprogramming, including the robust upregulation of the calcium-activated chloride channel ANO1. Both RNA-sequencing and RT-qPCR analyses confirm that ANO1 is among the most significantly elevated host genes in vGPCR-expressing HMEC-1 cells **(Figure 1B and 1E)**, alongside previously known vGPCR targets such as COX-2 and CXCL8. Our findings align with reports that viral manipulation of endothelial cells in diseases such as Kaposi’s Sarcoma involves the induction of pro-angiogenic and inflammatory genes and the suppression of vascular homeostasis signatures. [13,61–63]. Importantly, ANO1 upregulation was also observed in iSLK.BAC16 cells, a model of KSHV reactivation **(Figure 3B and 3D)**, indicating that ANO1 induction is not simply an artifact of vGPCR-overexpression. This convergence highlights that the vGPCR HMEC-1 model provides a useful framework in identifying the host factors that KSHV may modulate during infection.

Functional studies reveal that ANO1 is a critical effector of vGPCR-mediated cell survival under nutrient-limiting conditions. While ANO1 depletion did not affect viability in nutrient-rich conditions, serum starvation revealed a strict dependence on ANO1 for vGPCR to suppress apoptosis **(Figure 2B).** Rescue of ANO1 knockdown-induced cell death by the pan-caspase inhibitor Z-VAD-FMK demonstrates that the loss of viability is caspase-dependent, consistent with the reported anti-apoptotic role of ANO1 in cancers [18,24]. These findings suggest that ANO1 acts downstream of vGPCR to inhibit apoptotic pathways during cellular stress, potentially contributing to the apoptosis-resistant phenotype characteristic of KSHV-infected endothelial cells [28,29].

Interestingly, knockdown or pharmacological inhibition of ANO1 in KSHV-infected iSLK.BAC16 cells not only enhanced apoptotic cell death **(Figure 4)** but also increased lytic gene expression **(Figure 5)** and the production of extracellular infectious virions **(Figure 6).** Together, these results establish that ANO1 is required to suppress late-stage apoptosis during lytic KSHV infection, supporting a role for this calcium-activated chloride channel in sustaining host cell survival to optimize viral replication, while its loss appears to directly or indirectly favor viral gene expression and virion production, implicating ANO1 as a key modulator of host–virus interactions. The observed increase in viral titers following ANO1 loss could be related to apoptosis-associated membrane remodeling or vesicle release mechanisms that facilitate virion release [64,65]. Such coupling between apoptosis regulation and viral replication strategy has been documented in other herpesviruses and may represent a critical node in KSHV pathogenesis. In KSHV, one mechanism involves the latency-associated nuclear antigen (LANA) restraining p53-dependent apoptosis, thereby preserving the survival of infected cells [66]. This strategy parallels our observation that vGPCR-induced ANO1 upregulation in endothelial and KSHV-infected cells inhibits apoptosis, suggesting that ANO1 may be an additional host target co-opted to prolong cell viability during infection. Conversely, when apoptotic control is lost, such as with ANO1 depletion, cell death pathways can be activated in a way that facilitates viral release. Consistent with this, [67] demonstrated that apoptotic caspases can directly trigger viral replication and egress, supporting the idea that apoptosis, under certain conditions, enhances viral spread [67]. In our study, loss of ANO1 likely creates a pro-apoptotic environment that accelerates virus production. Additionally, KSHV v-cyclin binds CDK6 to promote cell cycle progression but also causes phosphorylation that inactivates cellular Bcl-2, promoting apoptosis. The viral Bcl-2 homolog protects against this effect, highlighting how KSHV balances apoptosis to benefit its life cycle [68]. Together, these findings position ANO1-mediated apoptosis inhibition within a broader network of KSHV strategies to fine-tune host survival, balancing latency and lytic reactivation for optimal replication dynamics.

Previous studies report that ANO1 is highly expressed in various types of cancers [17,29,59,69,70]. Knockdown of ANO1 expression in cancer cells leads to a significant amount of death, indicating that ANO1 expression is crucial in promoting cancer cell viability [18,70]. A recent study shows that ANO1 expression can modulate the activation of the Akt pathway, a classical pathway that controls a plethora of host physiological processes like apoptosis, metabolism, and cell survival. Not only can Akt activation lead to the inhibition of apoptosis through the activation of the nuclear factor kappa light chain enhancer of activated B cells (NF-kB), but it can also induce cell survival pathways by activating serum/glucocorticoid-regulated kinase (SGK) [18,71–73]. Furthermore, they show that the loss of ANO1 expression led to Akt inactivation and increased cell death, indicating that the activation of Akt is key to ANO1-mediated inhibition of host cell death [18]. Previous studies show that KSHV and vGPCR downstream signaling can also lead to the activation of the Akt pathway and NF-kB, which inhibits host cell apoptosis [28,74]. Another group demonstrated that KSHV-driven tumorigenesis is associated with pronounced epigenetic reprogramming, including hypo-methylation and upregulation of oncogenic pathways such as PI3K-Akt signaling, underscoring Akt pathway activation as a key feature of KSHV-induced sarcomagenesis [75]. Given these prior observations, our hypothesis is that vGPCR upregulates ANO1, and ANO1 contributes to the maintenance of PI3K-Akt signaling, thereby suppressing caspase-dependent apoptosis under stress. Loss of ANO1 would therefore reduce Akt signaling (or its downstream anti-apoptotic outputs), lowering the apoptotic threshold and allowing caspase activation during serum starvation or during KSHV lytic reactivation.

Although studies have shown that vGPCR-mediated inhibition of cell death is dependent on the activation of the Akt pathway, the interplay between ANO1 and the Akt pathway during the KSHV lytic phase is currently unknown [28]. Future experiments should determine the role of ANO1 in the activation of the Akt pathway during the KSHV lytic phase. The consistent upregulation of ANO1 in both HMEC-1 and iSLK.BAC16 transcriptomes point to a potentially broader role in KSHV biology beyond anti-apoptotic signaling. ANO1 is a calcium-activated chloride channel whose function directly intersects with calcium homeostasis [76]. Given that vGPCR signaling activates PLCβ, leading to IP3-mediated calcium release [15,28], and that calcium signaling influences both apoptotic cascades [77] and viral reactivation in herpesviruses [78], ANO1 may serve as a connecting pathway linking vGPCR-driven calcium fluxes to downstream survival and reactivation pathways. The enrichment of calcium signaling-related GO terms in our iSLK.BAC16 dataset further supports this connection **(Supplementary Figure 2B).**

Future studies should investigate the mechanistic interplay between ANO1 activity, calcium signaling, and caspase regulation in KSHV-infected endothelial cells. Pharmacological modulation of calcium flux and genetic dissection of downstream calcium-dependent kinases or phosphatases could clarify whether the ANO1 anti-apoptotic effect depends on calcium-dependent survival signaling or simply on chloride conductance. Similarly, examining specific caspase family members involved in ANO1-dependent survival will be critical to determining whether this pathway converges on intrinsic mitochondrial apoptosis or extrinsic death receptor pathways.

In conclusion, our findings identify ANO1 as a key vGPCR-induced host factor that supports endothelial cell survival under stress and limits KSHV lytic reactivation. By integrating transcriptional profiling with functional assays, we reveal a previously unrecognized role for ANO1 in KSHV biology and highlight calcium signaling and caspase-dependent apoptosis as promising avenues for therapeutic intervention. Given the ability of ANO1 inhibitors to both induce apoptosis and enhance viral reactivation, careful consideration of the timing and context of targeting ANO1 will be essential for translating these findings into antiviral or anti-tumor strategies.

## MATERIALS AND METHODS

### Plasmid Construction

To express SSF-tagged vGPCR in HMEC-1, we generated the plasmid pcDNA3.1(+)-SSF-vGPCR (6453 bp). The pcDNA3.1(+) backbone contains the CMV enhancer and CMV promoter for high-level expression, followed by a T7 promoter and a multiple cloning site (MCS). An N-terminal SSF tag (signal sequence + FLAG epitope) was inserted into the MCS region. Immediately downstream, the full-length vGPCR/ORF74 coding sequence was cloned in frame with the SSF tag, producing an N-terminally tagged SSF-vGPCR fusion protein. The insert region contains two components, SSF tag (78 bp) and vGPCR ORF (1029 bp), together making the SSF-vGPCR coding region 1107 bp, with additional flanking homology sequences included during synthesis to enable cloning. The plasmid retains all regulatory elements of pcDNA3.1(+), including the bovine growth hormone polyadenylation signal (bGH pA), SV40 promoter and origin, and AmpR and NeoR/KanR cassettes for bacterial and mammalian selection. Restriction sites such as ClaI, XhoI, BmtI, EcoRI, and XcmI flank the SSF-vGPCR region and were used for cloning and diagnostic digestion.

pcDNA3.1(+) (5428 bp) served as the Empty Vector control, containing the CMV promoter and standard expression elements without any inserted gene.

### Cell Culture

HMEC-1 were cultured in MCDB 131 media (Corning, 15-100-CV) supplemented with 10% fetal bovine serum (FBS) (Corning, 35-011-CV), 1ug/mL Hydrocortisone (Stem Cell Tech, 74142), 10ng/mL epidermal growth factor (Biovision, 4022-500), and 10 mM L-Glutamine (Gibco, 25030081) at 37°C and 5% CO2.

iSLK.BAC16 cells were cultured in DMEM media (Gibco, 1195-065) supplemented with 10% FBS (Corning, 30-249-CR), 10 mM L-Glutamine (Gibco, 25030081), (1% pen strep (Cytiva, SV30010), 1ug/mL puromycin (Mirus, 22103916), 250ug/mL G-418 (Fisher Bioreagents, BP673-5) and 1000ug/mL hygromycin (Corning, 30-240-CR)[4,35]at 37°C and 5% CO2. iSLK cells were cultured in similar media and conditions, but without hygromycin.

### Transient Transfection

Cells were seeded and grown overnight. HMEC-1 cells were transiently transfected to express N-terminal Signal Sequence FLAG-tagged vGPCR or an empty vector control to determine the role of vGPCR in host cell death and transcription. Transfections were performed using the TransIT-X2 Dynamic Delivery System (Mirus, MIR 6000) according to the manufacturer’s protocol. Cells were transfected for 48 hours, and then collected for downstream analysis. All downstream experiments used the generated signal sequence Flag-tagged viral G protein-coupled receptor, pcDNA3.1(+) _SSF-vGPCR, expression construct, and pcDNA3.1(+)_SSF-EV as a control.

### Virus and Induced Reactivation

iSLK.BAC16 cells stably maintain the KSHV genome and can be activated to enter lytic replication upon doxycycline and sodium butyrate treatment. iSLK cells do not contain the KSHV genome and are used as a control in our experiments. The iSLK.BAC16 cell line is latently infected with recombinant KSHV BAC16 encoding constitutive expression of EGFP. These cells encode the viral Replication and Transcription Activator (RTA) transgene, which is doxycycline (Dox) and sodium butyrate (NaB) inducible; this triggers the transition in iSLK.BAC16 from latent to lytic infection.

### RNA Extraction and RT-qPCR

HMEC-1 cells were seeded for bulk RNA sample preparation and other analyses. Transfections were performed at ∼70% as described previously. Cells were washed with PBS once, and RNA was isolated using the PureLink RNA Extraction kit (Invitrogen, 12183025) according to the manufacturer’s protocol. RNA was eluted in RNAse-free water, and the concentration of each sample was determined via UV-Vis Spectrophotometer. Two-step quantitative real-time reverse transcription-PCR (Bio-Rad) was used to measure expression levels of vGPCR and other known targets of vGPCR before sending the samples for sequencing. Relative levels of each transcript were normalized using the delta threshold cycle method to the abundance of GAPDH mRNA.

iSLK.BAC16 and iSLK cells were seeded for bulk RNA sequencing and other analyses. The next day, cells (∼70% confluency) were treated with Doxycycline (Dox) (1ug/mL) and Sodium Butyrate (NaB) (1ug/mL) for 48 h to induce lytic reactivation. After 48 h of Dox and NaB induction, iSLK/iSLK.BAC16 (+/- Dox) cells were washed with PBS once, and RNA was extracted using the PureLinkTM RNA Mini Kit according to the manufacturer’s protocol. Two-step quantitative real-time reverse transcription-PCR (Bio-Rad) was used to measure expression levels of vGPCR and other lytic genes for reactivation. Relative levels of each transcript were normalized using the delta threshold cycle method to the abundance of Tubulin mRNA.

cDNA was generated from RNA using the iScript cDNA Synthesis Kit (Bio-Rad, 1708841). Briefly, 1 μg of RNA was used per reaction primed. The reverse transcription reaction was performed according to the manufacturer’s protocol. qRT-PCR was performed using a QuantStudio 3 Real-Time PCR Detection System instrument (Applied Biosystems). cDNA generated from extracted RNA was used as the input for the qRT-PCR, with amplification by the iTaq Universal SYBR Green Supermix (Bio-Rad, 1725124). All samples were tested in triplicate or more. All primers were synthesized by Integrated DNA Technologies, Inc. (San Diego, CA), and their sequences are described in Supplementary Table 1.

### RNA-sequencing

HMEC-1 cells were seeded in 10 cm dishes for bulk RNA sample preparation. Transfections were performed at ∼70% confluency, and RNA was extracted after 48 hours of transfection as described previously. RNA was sent to NovoGene (CA, USA) for sequencing. Messenger RNA was purified from total RNA using poly-T oligo-attached magnetic beads. After fragmentation, the first strand cDNA was synthesized using random hexamer primers, followed by the second strand cDNA synthesis using either dUTP for a directional library or dTTP for a non-directional library. For the non-directional library, it was ready after end repair, A-tailing, adapter ligation, size selection, amplification, and purification. For the directional library, it was ready after end repair, A-tailing, adapter ligation, size selection, USER enzyme digestion, amplification, and purification. The library was checked with Qubit and real-time PCR for quantification and a bioanalyzer for size distribution detection. Quantified libraries were pooled and sequenced on Illumina platforms according to effective library concentration and data amount. The clustering of the index-coded samples was performed according to the manufacturer’s instructions. After cluster generation, the library preparations were sequenced on an Illumina platform, and 150-bp paired-end reads were generated.

iSLK and iSLK.BAC16 cells were seeded in 6-well plates, grown overnight, induced using Dox and NaB, and RNA was extracted as described previously. RNA samples were purified using DNase 1 Amplification Grade (Invitrogen, 18068015) before sending the samples for sequencing at the UT Dallas Genomic Core. Total RNAs were purified and subjected to Illumina stranded mRNA library preparation according to the manufacturer’s instructions (Illumina). The libraries were quantified by Qubit (Invitrogen) and the average size of the libraries was determined using the High Sensitivity NGS fragment analysis kit on the Fragment Analyzer (Agilent). The normalized libraries were then sequenced on an Illumina NextSeq2000 sequencing platform with 100-bp paired-end reads.

### RNA-seq Data Analysis

#### Quality control

Raw data (raw reads) of fastq format were first processed through in-house Perl scripts. In this step, clean data (clean reads) were obtained by removing reads containing adapters, reads containing poly-N, and low-quality reads from the raw data. At the same time, Q20, Q30, and GC content of the clean data were calculated. All the downstream analyses were based on clean data with high quality.

#### Reads mapping to the reference genome

The human reference genome, GRCh38, and its corresponding annotations, were obtained from NCBI Datasets and indexed using the software Spliced Transcripts Alignment to a Reference (STAR) v2.7.11b. Paired-end reads were converted from the BCL file format to the fastq file format using Illumina’s bcl2fastq v2.20 and filtered based on quality before being aligned to the indexed genome using STAR v2.7.11b to produce Bam files.

#### Quantification of gene expression level

The ‘featureCounts’ function from the Rsubread v2.20 R package was used to count the number of reads mapped to each gene for each Bam file. These counts were normalized to gene length and sequencing depth, calculating the expected number of fragments per kilobase sequenced per million base pairs sequenced (FKPM), using the ‘fkpm’ function from the DESeq2 v1.48.0 R package.

#### Differential expression analysis

Differential expression analysis of two conditions/groups (two independent biological replicates per condition for the HMECs and five independent biological replicates per condition for the iSLKs) was performed using the DESeq2 R package (1.48.0). DESeq2 determines statistical significance of differential expression through fitting the transcript counts of each gene to a general linear model (GLM) using negative binomial regression. The resulting *p*-values were adjusted using Benjamini and Hochberg’s approach for controlling the false discovery rate. Genes with an adjusted *p*-value <=0.05, Log2FoldChange < -1 for HMEC-1 and adjusted *p*-value <=0.05, Log2FoldChange < -2 iSLK cells found by DESeq2 were assigned as differentially expressed. R and R studio scripts were used to create the heatmap and volcano plot, respectively, to visualize the DE genes.

### siRNA Transfections and Serum Starvation

For siRNA transfections, cells were seeded in a 6-well plate at a density of 1.65 x105 cells/ well or 50% confluency, then transfected with either pcDNA3.1(+)_SSF-EV or pcDNA3.1(+) _SSF-vGPCR plasmids using TransIT-X2 Dynamic Delivery System (Mirus, MIR 6000) according to the manufacturer’s protocol. At 24-hr post-transfection, cells were transfected with siANO1 (Qiagen, Germantown, MD) or non-targeting scrambled control All-Star Negative siRNA AF 488 (siASN) (Qiagen, 1027292) at a final concentration of 25 nM siRNA, using TransIT-X2 Dynamic Delivery System (Mirus, MIR 6000) according to the manufacturer’s protocol. 36-hr post-siRNA transfections, media will be replaced with either supplemented or serum-deprived DMEM media (only L-Glut added). After 12 hours of serum deprivation, trypan blue cell viability assay, apoptosis rescue experiment, and flow cytometry were performed as described.

iSLK.BAC16s and iSLK cells were seeded in a 6-well plate at a density of 1.65 x105 cells/ well or 50% confluency. The next day, cells were transfected with siANO1 (Qiagen, Germantown, MD) or non-targeting scrambled control All-Star Negative siRNA AF 488 (siASN) (Qiagen, 1027292) at a final concentration of 25 pmol siRNA, using Lipofectamine RNAiMAX (Mirus, MIR 6000) according to the manufacturer’s protocol. 24-hour post-transfection, cells were treated with -/+ Dox/NaB to induce reactivation. 36-hr post-lytic induction, media will be replaced with either supplemented or serum-deprived DMEM media (only L-Glut added). After 12 hours of serum deprivation, a trypan blue cell viability assay, an apoptosis rescue experiment, and flow cytometry with Apotracker-Alexa Fluor 647 will be performed as described.

The knockdown of ANO1 was accomplished by FlexiTube GeneSolution siRNA (Qiagen, 1027416) according to the manufacturer’s protocol, with 4 types of siRNA for HMEC-1 cells, and only TMEM16A_6 ANO1 knockdown efficiency for BAC16 cells was confirmed using RT-qPCR analysis before moving forward with the cell viability assays.

### Trypan Blue Cell Viability Assay

Cells were seeded and treated as described in the above section. Cell media was collected in different tubes, cells were washed once with PBS, trypsinized (Gibco, 25200056), neutralized with fresh media, which were then collected in the same tubes. 0.4% Trypan Blue (Life Technologies, T10282) was mixed with aliquots of collected cells at a 1:1 ratio. Then the suspension was loaded onto a cell counting slide (Bio-Rad, 1450015) and cells were counted using the Bio-Rad TC-20 automated cell counter. Alternatively, cells were also counted using the classic hemocytometer and scored with the VWR trinocular Inverted microscope for confirmation.

Viable and dead cells were counted to obtain the total cell count and live/dead cell percentage. Cell death was calculated using the formula below.

Cell death percentage=[(Total # dead cells)/(Total # live cells + Total # dead cells)]*100

### Apoptosis Rescue Experiment

To determine whether cell death was due to apoptosis, a rescue experiment was performed using the Z-Vad pan-caspase inhibitor (Z-VAD-FMK) (Promega, G7232). Cells were seeded and treated as described in the "Serum-Starved Conditions and siRNA” section above. 36-hr post-transfection, the media was replaced with either full serum, serum-free media, or serum-free media containing Z-VAD-FMK at a final concentration of 25 µM to inhibit caspase activity or an equivalent volume of DMSO as a control. The Trypan blue cell viability assay was done as previously described.

### Flow Cytometry

HMEC-1 and iSLK.BAC16 cells were seeded and treated as described above. Media was collected, cells were washed once with PBS, and trypsinized and pelleted by centrifugation at 300g for 5 minutes. eBioscience™ Annexin V Apoptosis Detection Kit (Invitrogen 88-8007-74, 88-8005-74) was used to detect early and late apoptotic cells. The supernatant was discarded, and the cell pellet was washed once with PBS and once with 1X Binding buffer provided in the kit. Cells were then resuspended in 100 μL 1X Binding buffer. 5 μL of Annexin V-FITC/APC was added to the resuspended cells and incubated for 10-15 minutes at RT. Cells were then washed in 1X Binding buffer and then resuspended in 200 μL 1X Binding buffer. 5 μL of Propidium Iodide (PI) was added to the resuspended cells. For the flow cytometry analysis, after staining, the samples were analyzed within 30 minutes - 1 hour using a flow cytometer equipped with filters suitable for detecting FITC (Excitation/Emission 490/525 nm) APC (Excitation/Emission 650/660nm) and PE-Texas Red (Excitation/Emission 535/617 nm) fluorescent dyes. A BD Fortessa SORP cytometer, available at the UT Dallas Biological Sciences core, was used for FACS analysis.

During data collection, a minimum of 10,000 events per sample were recorded to ensure statistical significance. Acquired flow cytometry data were analyzed using FlowJo v11.0.2, BD FACSDiva software. Early and late apoptotic populations were identified by gating on forward scatter (FSC) and side scatter (SSC) to exclude debris and doublets. Quadrant gating differentiated live, early, early-late, and late-apoptotic cell populations. Annexin-V^+^/PI^-^ cells (Quadrant Q1) were classified early-apoptotic, Annexin V^+^/PI^+^ cells were classified early-late apoptotic(Quadrant Q2), Annexin-V^-^/PI^+^ cells (Quadrant Q3) were classified late-apoptotic, Annexin-V^-^/PI^-^ cells (Quadrant Q4) were classified as viable. Data were exported, and

percentages of each population were normalized to non-reactivated controls and quantified for statistical analysis using GraphPad Prism 8.0.1 software. All experiments were performed at least in biological triplicate, with statistical significance determined using one-way ANOVA.

### Drug Treatment

iSLK.BAC16 cells were seeded in a 6-well plate at a density of 1.65 x105 cells/ well or 50% confluency. The next day, cells were treated with different concentrations of the ANO1-inhibiting drugs, Ani9 (TOCRIS, 6076) and Diethylstilbestrol (DES) (Sigma-Aldrich, 46207). At 24-hr post-treatment, cells were treated with -/+Dox/NaB to induce lytic reactivation. At 48-hr post-lytic reactivation, supernatant was collected and saved to perform titer experiments. The cells were harvested for qPCR analysis or used for flow cytometry analysis.

### Extracellular Virus Titers

All iSLK.BAC16 cell supernatants were collected post-experimental treatment and subjected to centrifugation to clear debris at 300g for 10 minutes. iSLK cells were seeded in 12-well plates at 70% confluency 24 h before infection. 500ul of supernatant supplemented with polybrene (10ug/ml) from treated cells was added to iSLK cells.

Infection was enhanced by spinoculation at 300g, 37 °C for 1 h (Acceleration 7, Deceleration 7) and then incubated for 4 hr at 37 °C with interval shaking every 30 mins. Infectious medium was replaced with fresh iSLK medium after 4h. Titers were imaged after 48 h on ZOE (BIO-RAD ZOE Fluorescent Cell Imager) for GFP+ cells, images were quantified using ImageJ software, and stats were performed.

### Statistical Analysis

Figures and statistical analyses were generated using GraphPad Prism 8.0.1 software. Experiments were performed at least in triplicate. Data are presented as the mean ± SEM. Experimental conditions were compared to control conditions using a t-test or one-way ANOVA. Significant p-values are noted in each figure (* ≤ 0.05; ** ≤ 0.01; *** ≤ 0.001; **** ≤ 0.0001), ns=non-significant change.

## Acknowledgements

We would like to thank both San Francisco State University as well as The University of Texas at Dallas for their support on this project. We would also like to thank all members of the Sanchez lab for their creative input throughout this study. We acknowledge the Genome Center in UTD Core Facilities for mRNA library preparation and sequencing. We would further like to acknowledge and thank the Biology Department at the University of Texas at Dallas for helpful discussions on this project.

## Supplementary Information

### Supplementary Figures

**Supplementary Figure 1. vGPCR overexpression leads to global changes in the host cell transcriptome. (A)** Heatmap showing all of the differentially expressed cellular transcripts, including 6 lncRNAs in vGPCR-transfected cells compared to EV controls. **(D)** Gene Ontology presenting the top 10 most significantly activated pathways and the top 10 most significantly suppressed pathways. **(C)** RT-qPCR validation of some of the top upregulated genes, previously studied in vGPCR-dependent transcriptional alterations. Generated in GraphPad. Data represent mean ± SEM from [n=3] independent experiments. Statistical significance was determined by t-test; ***, *P* ≤ 0.001; *, *P* ≤ 0.1. Genes with adjusted *p*-values 0.05 and Log2FoldChange -1, 1 were plotted. EV, Empty Vector.

**Supplementary Figure 2. Global transcriptomic changes in iSLK.BAC16 cells during KSHV reactivation. (A)** Heatmap visualization of the top 50 upregulated genes across replicates in iSLK.BAC16 cells **(B)** Gene Ontology presenting the top 10 most significantly activated pathways and top 10 most significantly suppressed pathways in iSLK.BAC16 non-reactivated versus DOX/NaB reactivated iSLK.BAC16 cells.

**Supplementary Figure 3. Knockdown efficiency of ANO1 in HMEC-1 and BAC16 cells. (A)** The knockdown efficiency of ANO1 in vGPCR-transfected HMEC-1 cells is observed to be ∼70%. Our RNA-sequencing data show little to no ANO1 expression in endothelial cells prior to vGPCR overexpression (normalized FPKM value for ANO1 for EV is 2.04 compared to 41.47 for vGPCR-transfected cells). **(B)** The knockdown efficiency of ANO1 in BAC16 cells is observed to be ∼70%. Generated in GraphPad. Data represent mean ± SEM from [n ≥ 3] independent experiments. Statistical significance was determined by t-test and one-way ANOVA; ****, *P* ≤ 0.0001. ASN=All-Star Negative; D/N=Doxycycline/Sodium Butyrate.

**Supplementary Figure 4.** K**i**ll **curve analysis of ANO1 inhibitors in iSLK.BAC16 cells.** (A) iSLK.BAC16 cells were treated with increasing concentrations of DES (0-15 μM) or DMSO control, and cell death was quantified by trypan blue exclusion assay. DES concentrations used in subsequent experiments (≤10 μM) did not induce significant cell death, whereas higher concentrations resulted in increased toxicity. (B) iSLK.BAC16 cells were treated with increasing concentrations of the selective ANO1 inhibitor Ani9 (0-45 μM) or DMSO control, and cell death was assessed by trypan blue exclusion assay. Ani9 concentrations used in downstream experiments (≤30 μM) did not significantly affect cell viability, while higher doses induced increased cell death. Data are presented as mean ± SEM from independent experiments (n ≥ 3). Statistical significance was determined using one-way ANOVA; ***p < 0.001; **p < 0.01; *p < 0.05; ns, not significant.

**Supplementary Table 1:** RT-qPCR Primer Sequences

